# Spatio-temporal modelling for the evaluation of an altered Indian saline Ramsar site and its drivers for ecosystem management and restoration

**DOI:** 10.1101/2021.03.02.433535

**Authors:** Rajashree Naik, L.K. Sharma

## Abstract

Saline wetlands are keystone ecosystems in arid and semi-arid landscapes that are currently under severe threat. This study conducted spatio-temporal modelling of the largest saline Ramsar site of India, in Sambhar wetland from 1963-2059. One CORONA aerial photograph of 1963 and Landsat images of 1972, 1981, 1992, 2009, and 2019 were acquired and classified under 8 classes as Aravalli, barren land, saline soil, salt crust, saltpans, waterbody, settlement, and vegetation for spatial modelling integrated with bird census, soil-water parameters, GPS locations, and photographs. Past decadal area statistics state reduction of waterbody from 30.7 to 3.4% at constant rate (4.23%) to saline soil. Saline soil increased from 12.4 to 21.7% and saline soil converted to barren land from 45.4 to 49.6%; saltpans from 7.4 to 14% and settlement from increased 0.1 to 1.3% till 2019. Future predictions hint at a net increase of 20% by wetland, vegetation by 30%, settlement by 40%, saltpan by 10%, barren land by 5%, and net loss of 20%, each by Aravalli and salt crust. The biggest loss of 120% was seen by saline soil converted to barren land. Notably, 40% of the current wetland will be lost by 2059. Additionally, soil-water parameters result state a loss of saline character of wetland ecosystem; subsequently bird statistics indicate a shift in migratory birds disturbing the wetland food web. India has been losing a critical habitat of migratory birds, halophytes, and halophiles, along with livelihood. This study looks to bridge the missing link from local to global wetland ecological disconnect, providing thereby lake management and restoration strategies.

## Introduction

Due to human influences, freshwater shortage and water stress are argued to be linked [1]. The anthropogenic alteration of natural resources has dramatically modified hydrological systems. [2]. Thus, considering anthropological influences, it is critical to conduct a systematic assessment of water resources in arid and semi-arid regions [3]. Saline lakes make up 44% and 23% of the volume and area of all lakes [4]. These occur mostly in arid and semi-arid regions as endorheic basins [5]. They have high ecological, economic, cultural, recreational, and scientific values [6]. However, anthropogenic activities, especially salinization, water inflow diversions, construction of hydrological structures, pollution, mining, biological disruptions, exotic species invasion have already threatened these lakes [7] resulting in changes in hydro-patterns, water budget, and hydrological communications, habitat alteration, loss of productivity and connectivity among wetland complexes [8]. Saline lakes around the world are predicted to suffer from extended dryness, reduced hydroperiod, increased salinity, or complete desiccation by 2025 as already seen in Aral Sea, Lake Urmia, Owens Lake, Tarim Basin, Salton Sea [4]. These directly affected the billion-dollar global markets of shrimp, mineral industry, and ecologic disruption [4].

According to the Government of India (GoI), India ranks third in global salt market after China and USA contributing approximately 230 million tons exporting to 198 countries. Major importers are Bangladesh, Japan, Indonesia, South and North Korea, Quatar, Malaysia, U.A.E, and Vietnam. In India, 96% of salt is produced from Gujarat, Tamil Nadu, and Rajasthan with 76.7%, 11.16%, and 9.86% respectively from sea, lake, sub-soil brine, and rock salt deposits. Rajasthan state exports 22.678 million tonnes of salt worth 340.17 billion dollar to global market extracted from inland lake brine in Nawa, Kuchhaman, Rajas, Phalodi, Sujangarh, and Sambhar lake [9]. This current study is in Sambhar Salt Lake (SSL), which is a gateway to the Thar Desert in India. It is the seventh Ramsar site of the country, designated on 23 March 1990, under criteria A with site No. 464 [10] and Important Bird Area No. IN073 [11]. Once it was a haven for 279 migratory and resident birds [12], which currently serves as a refuge for 31 migratory. Interestingly, despite its many years of corruption, it is not included in country’s protected network. The greatest threat to this lake is illegal encroachment in the core area, which nearly destroyed the lake’s identity [13]. Many illegal tube wells have been drilled, and long pumps are used for groundwater over-extraction. Prior encroachment has turned it into a large capital-intensive corporate business. [14]. It is hard to ignore illegal consequences even after repeated NGT intervention [15]. Importantly, SSL does require urgent restoration to its pristine condition.

Unfortunately, saline lakes in remote and inaccessible locations are little studied. Conventionally, numerous in-situ studies were conducted for saline lakes on phillipsite [16], chemical and biological properties [17], phytoplankton [18], primary productivity [19], stable isotopes [20], geochemistry [21]. In SSL, studies have been conducted for birds [22], halo-tolerant species identification by [23] and isolation [24], their characterization [25, 26]; on *Dunaliella sp.* [27] on lake formation [28]; on its limnological aspect [27]; on paleoclimatological conditions [26]; for sensor calibration and validation [29] and on extremophilic algal assessment [13]. These studies, however, are time-consuming, tedious, and expensive. Such research may not match fast-changing ecosystems.

Saline wetlands have dynamic hydro-periods [30]. These ecosystems usually experience Land Use Land Cover (LULC) modification seasonally [31]. So geospatial modelling is inevitable for these ecosystems. Popular models of contemporary literature are cellular-based and agent-based (or their hybrid) [32]. Cellular-based models consider both spatial and temporal components of LULC dynamics [33]. They are easy to standardize and competent to simulate wetland complexes [34]. These have been extensively used for past LULC trend analyses. However, future prediction enables the comprehension of sustainable management, restoration, combat desertification, biodiversity loss assessment and, water budgeting [35]. This research is conducted to complement the United Nations Decade on Ecosystem Restoration from 2021–2030 using aerial photograph of CORONA (1969), multispectral data sets, and ground information with Cellular Automata/Markov model to assess and predict the impact of salt pan encroachment under three-time frame as past (1969-2009), present (2019) and future (upto 2059).

## Material and methods

### Study area

SSL is in semi-arid climatic region of Rajasthan (Fig 1**Error! Reference source not found.**) with 26°52’to 27°Ü2’ N; 74°54’ – 75°14’ E running ENE–WSW direction in elliptical shape [11]. In 1961, the government acquired the region on a 99-year lease under the Ministry of Commerce and Industry [10]. SSL is 230 km^2^ (22.5 km in length and 3-1 km in width) [23]. One of the world’s oldest mountain range, Aravali surrounds it in the north, west, and south-east directions, extending up to 700 m [11]. Its maximum altitude is 360 m above mean sea level, with 10 cm per km slope. Ephemeral streams (Mendha, Rupnagar, Khandel, Kharian) form the catchment of 5,520 km^2^. Mendha in fact is the largest feeder river that originates in the north from Sikar district and drains out in 3600 km^2^· Notably, it experiences tropical climate, and its soil consists of silt and clay. Some part of the basin is calcareous, while most of it is argillaceous; it is rich in salts of sodium, potassium, calcium and magnesium cations and carbonate, bicarbonate, chloride, and sulfate anions. It appears white especially in areas with rich salt content; appears grey in areas with less salt content, and brown with no salt content [36]. Importantly, SSL at large, experiences distinct summer (March-June), rainy seasons (July-September), and winter (October-February). Overall, it receives around 500 mm of rainfall every year, while it enjoys 250-300 sunny days [11]. Additionally, the average temperature is about 24.4 °C, going up to 40.7 °C in summer, and below 11 °C in winter [36]. Further, during rainy seasons, it looks like a muddy blackish wetland [25]. It has almost 3 m depth during monsoons, but shallows down to 60 cm during dry periods. Except for reservoir and saltpans, the whole lake dries up exposing salt flakes during summer [37]. A 5.16 km long dam divides into two unequal parts (77 km^2^ towards east as reservoir and rest 113 km^2^ is wetland) [26]. It receives migratory birds of Central Asian, East Asian, and East African flies. Invertebrates, amphibians, crustaceans come through rivers during monsoon when salinity is low [23]. Moreover, it provides shelter to 37 herbs *(Portulaca oleracea, Salsola foetida, Suaeda fruticose)* 14 shrubs *(Salvadora oleoides, Salvadora persica, Sericotoma pauciflorum)* 14 trees *(Acacia nilotical, Acacia Senegal, Anogeissus pendula)* 15 grasses (*Apluda mutica, Aristida adscensionis, Cenchrus ciliaris)* 6 chlorophycea *(Chamydomonas sp.,Dunelialla salina, Oedogonium sp.)* 25 Cyanophyceae *(Lyngbya sp., Merismopedia sp., Microcole, us sp.)* and 7 Bacilariophyceae *(Cymbella sp., Melosira sp., Navicula sp)* species [38].

**Fig 1.**
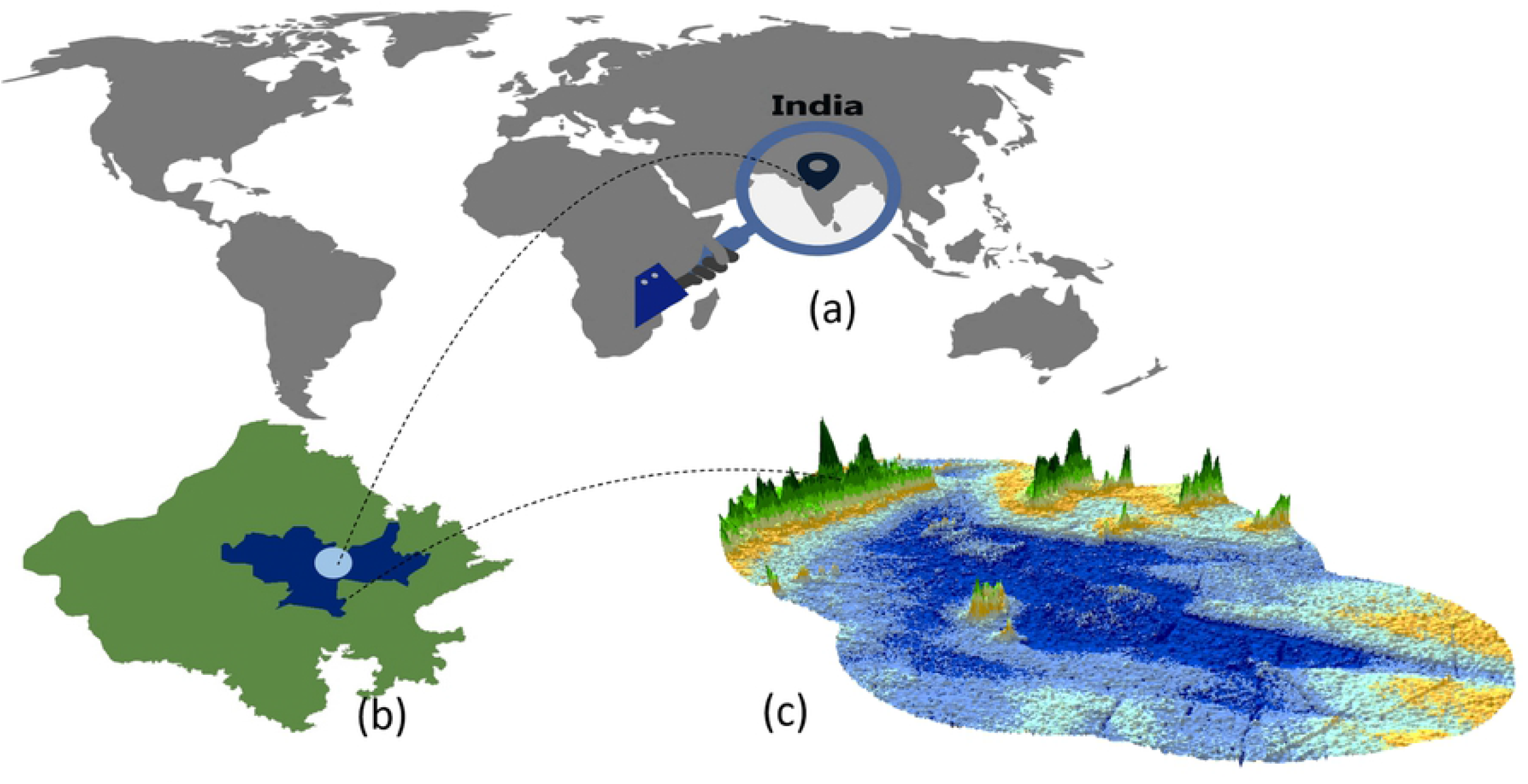
Study area, (a) Indian saline Ramsar sites (b) SSL amid Nagaur, Jaipur and Ajmer.

### Data processing

We were interested in knowing the lake’s status, especially in winter, when adequate surface water is expected and as mentioned earlier it naturally dries in summer. LULC classification on a decadal scale was carried out for past, current, and future changes. Notably, only one aerial image of CORONA was obtained, which is well before the start of any satellite programmes. This photograph has only been digitized for visual interpretation of LULC classes; however, being a declassified image, we could not calculate the area. Satellite data including Landsat-MSS (Multispectral Scanner System) of 16 November 1972 and 18 October 1981, Landsat-5 Thematic Mapper (TM) of 25 November 1992 and 8 November 2009, and Landsat-8 Operational Land Imager (OLI) of 20 January 2019 were also collected. Images of 1972 and 1981 have 60m spatial resolution, while the rest of the images have 30m resolution. The best available cloud-free images of Landsat satellite were downloaded within this season. Additionally, the study was not impacted by droughts or floods. The study years, 1972, 1981, 1992 received rainfall above 500 mm, while 2009 and 2019 received just about average rainfall [36]. The methodology followed has been shown in Fig 2.

**Fig 2.**
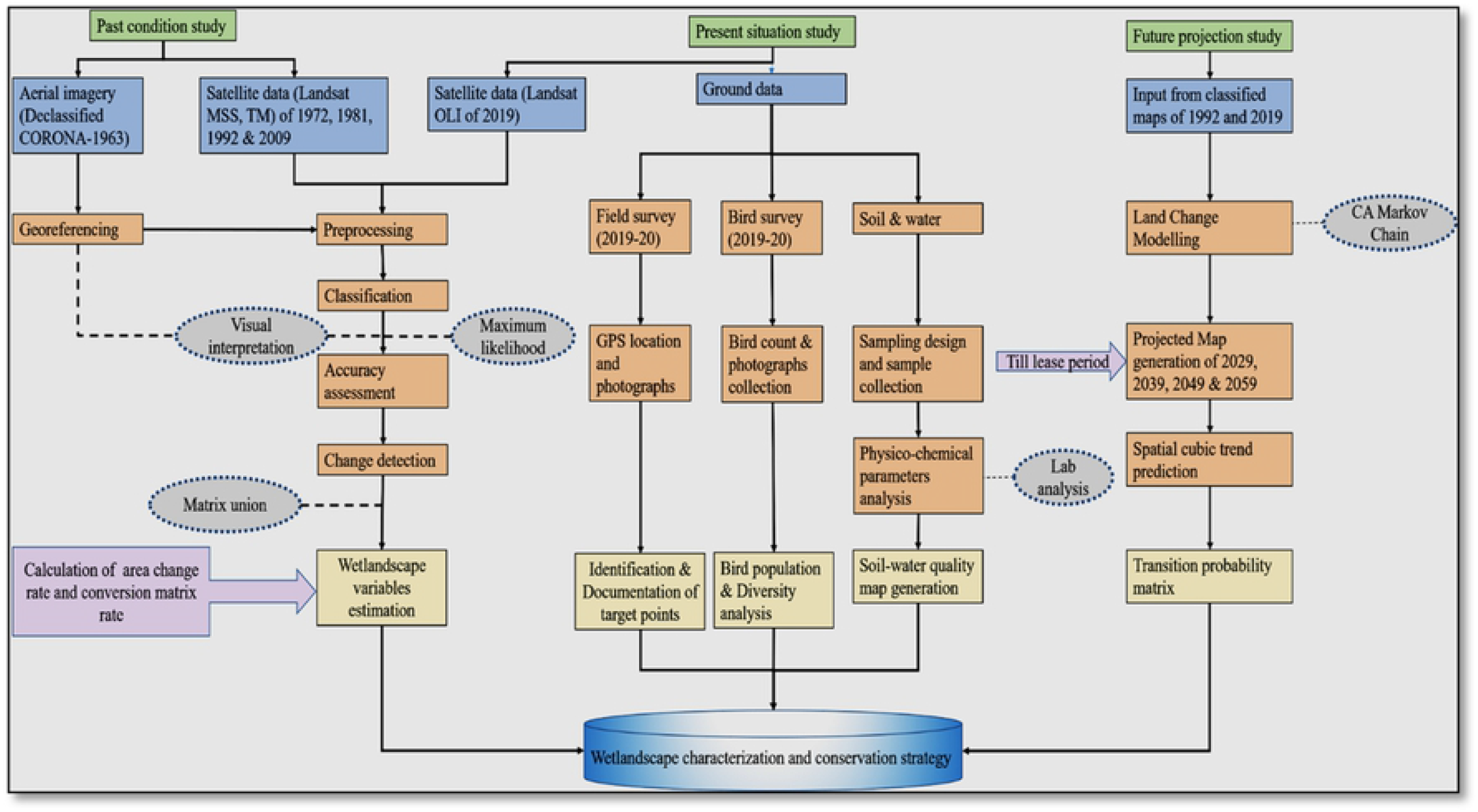
Methodology flowchart.

All of the downloaded images were geo-referenced and atmospherically and geometrically corrected. To maintain uniformity, the pan-sharpening of 1972 and 1981 images was done to 30m spatial resolution. Toposheet from Survey of India (1954) at 1:26,000 scale was used for boundary delineation. SSL was digitized and 3 km buffer was selected, as it is declared as an eco-sensitive zone, according to Rajasthan State Forest Department. For classification, pixel-based method was used using ERDAS Imagine, 2014, while the final maps were composed using Arc GIS 10.5. Further, SSL was divided into eight classes using supervised classification method; they include waterbody, saltpan, salt crust, vegetation, the Aravalli mountain range, saline soil, barren land, and settlement. The water bodies represent the wetland areas, which do not come under the reservoir. It appears dark-light blue in True Color Composite (TCC). Saltpans on the other hand, are the salt-producing units; salt crust represents high salt deposition area, appearing white, while vegetation appears green, and are occupied by both xerophytes and halophytes, Aravalli represents the hill ranges, saline soil represents the terrestrial part of the lake with both soil and salt content appearing grey, barren land represents area without salt content appearing brown, while settlement represents built-up area surrounding SSL. Moreover, past change detection was conducted for 47 years (i.e., 1972-2019) on a decadal scale. Notably, LULC of each image was estimated using pixel-based classification. Supervised Maximum Likelihood classification method (MLC) was applied. 69 GPS locations were obtained from in and around the lake during soil and water sample collections, bird census, and validation of classification shown in Fig 3.

**Fig 3.**
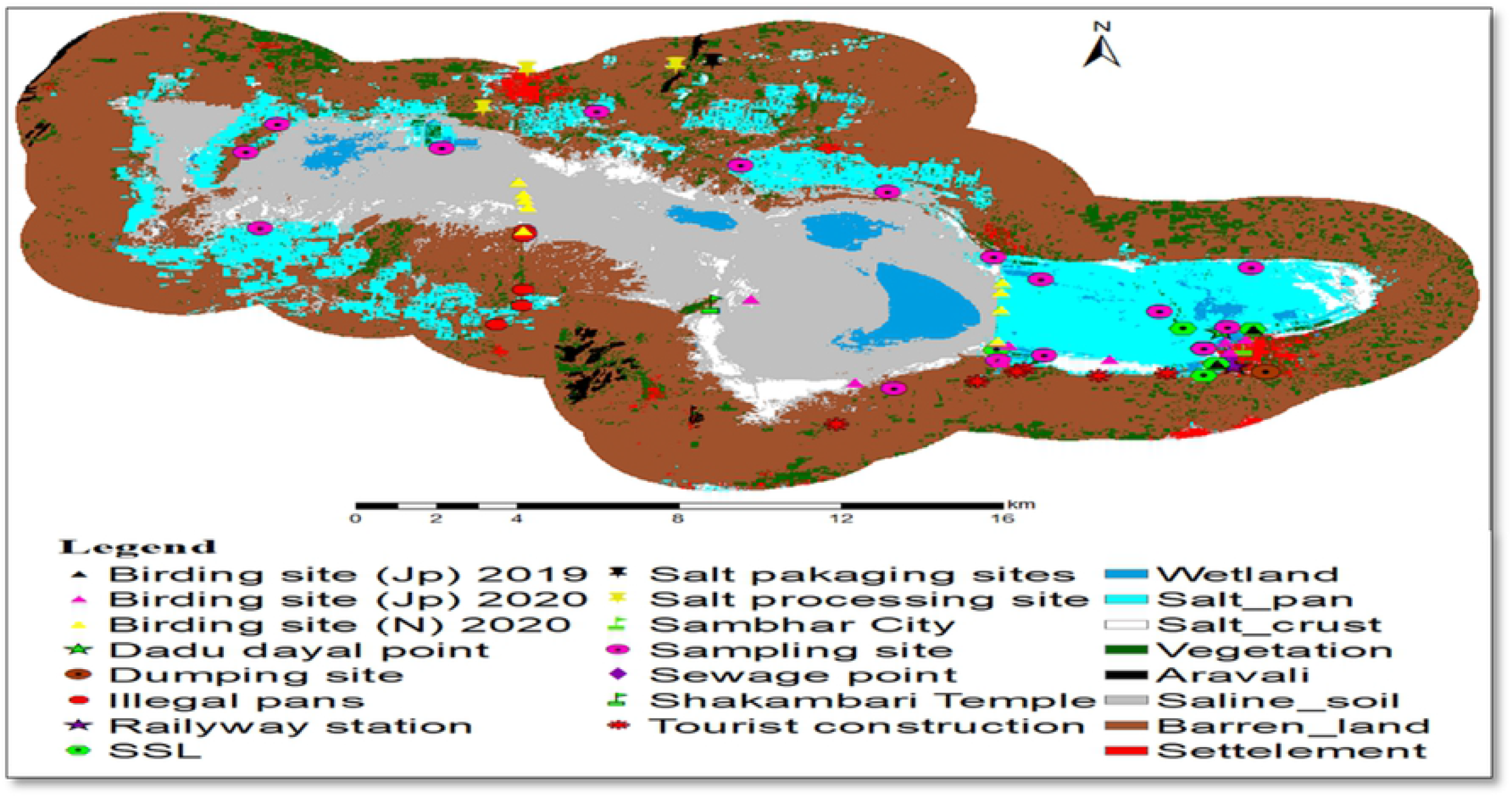
Location of field points.

Target locations were pre-defined for each class and sampling. Detailed field research were carried out on 13 February 2019 (winter); 10 April 2019 (summer); 30 June 2019 (monsoon), and 6 January 2020. Additionally, out of 69 GPS locations, 48 points were used for classifications, and other points for accuracy assessment. Primary landmarks like historical sites (Shakambari temple, Devyani Sarovar, Dadu Dayal point, Sambhar City, Railway Station), along with birding sites, dumping sites, illegal pans, sewage points, tourist construction sites, salt processing, and packaging sites were identified (Fig 4 (a-f)) and historical photographs were collected from [38] as given in Fig 4 (g-h). The Google Earth was used for accuracy assessment of the past images. Dynamic degree was calculated using [40]

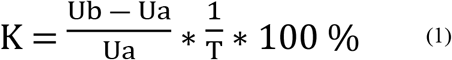

**Fig 4.**
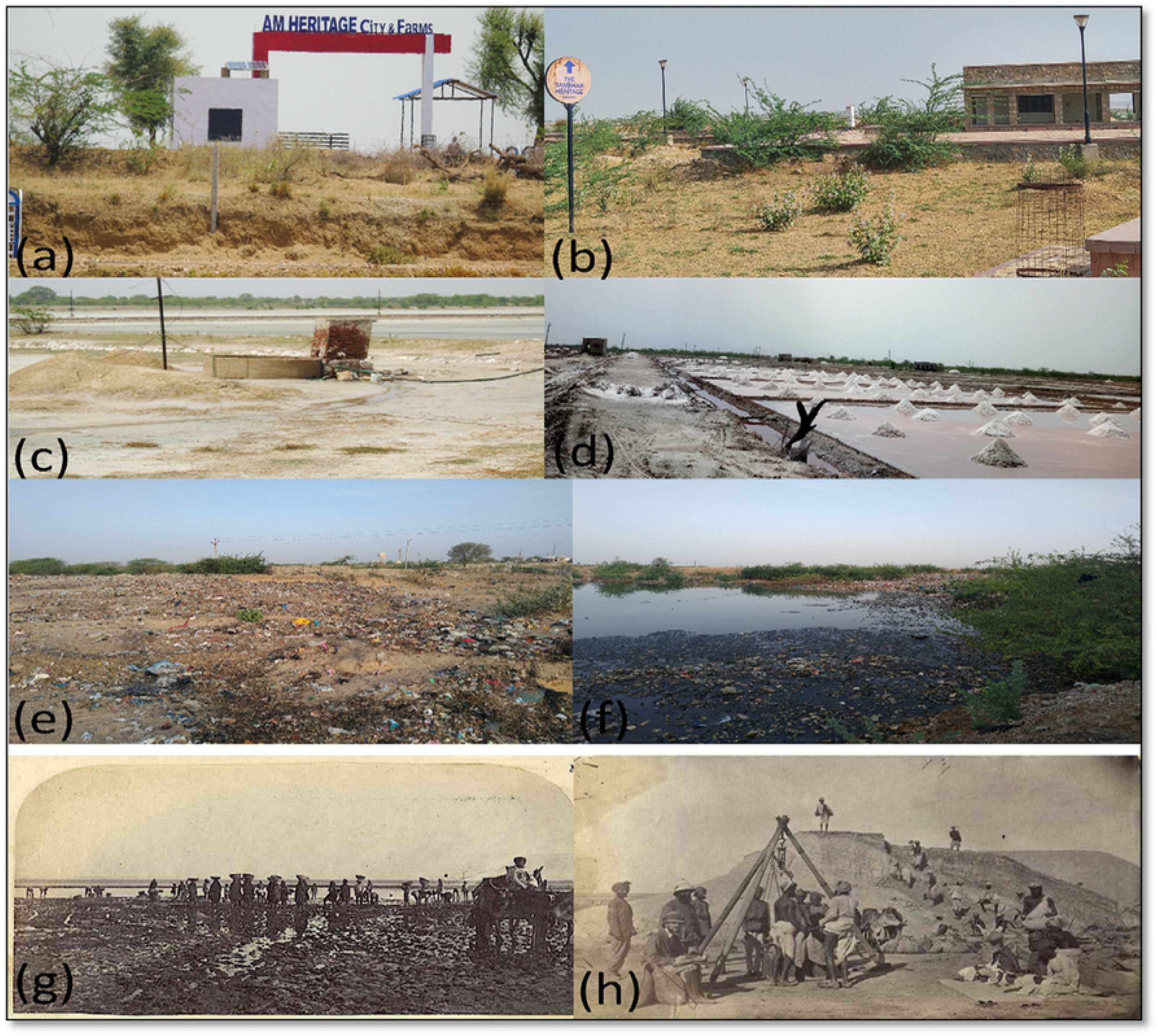
Field Photographs (a) & (b) construction sites, (c) & (d) surface wells, saltpans with illegal electrical cables (e) & (f) domestic pollution sources, and (g) & (h) historical photographs of lake and salt production.

**Fig 5.**
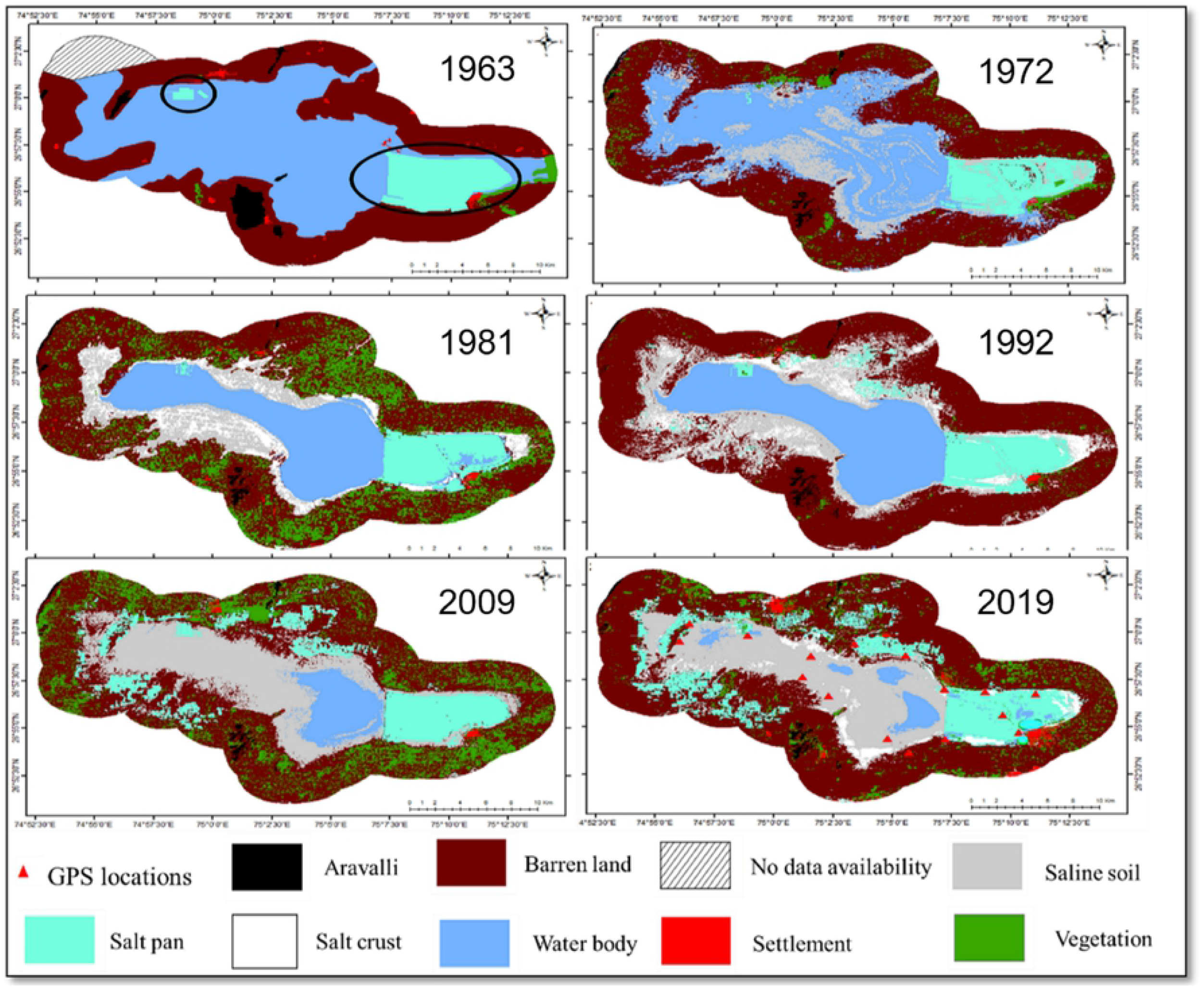
LULC maps 1963, 1972, 1981, 1992, 2009 and 2019 of SSL.

K is the land use dynamic degree, calculated as percent LULC change per year, both Ua and Ub represent areas under a specific annual LULC, while T represents the time in years. Dynamic matrix was generated between 1981-2019 to estimate LULC transfer. Notably, to quantify transition matrix thematically, equal number of classes are required. Classified image of 1981 was taken instead of 1972 for matrix analysis, as SSL was subject to flash floods caused by 1000 mm of rainfall in 1971. So, due to rising water levels, no salt crust was observed in the raw image. For future projection of four decades (i.e., from 2029-2059), Cellular Automata Markov Chain Model of Land Change modelling (LCM) was conducted using Terra Set software. Classified images of 1992, 2009, and 2019 (two decades representatives) were used for forecasting. The quantitative results were achieved using LCM, net change of each class was calculated and then the factors of change were calculated.

Moreover, to assess the current situation, three parameters (i.e., soil, water, and bird count) were chosen. Sixteen samples, each for soil and water were collected by stratified random sampling method. These collected samples were further analyzed in the laboratory of Department of Environmental Science as per American Public Health Associations (APHA) guidelines. pH and electrical conductivity were examined using respective electrode. Other parameters like salinity, chloride, carbonate, total organic carbon of soil and total dissolved solid, salinity, hardness, carbonate of samples was analyzed using titration method.

Bird censuses were conducted on 11-Jan-2019, and 6th and 7th January 2020. From the survey, 29 and 32 bird species with total of 1124 and 43,445 bird count were recorded in the respective years (Table 1). Good rainfall increased bird counts in 2019. 10 bird species like Black Crowned Night Heron, Greater Flamingo, Lesser Flamingo, Gadwall, Little Ringed Plover, Kentish Plover, Red Wattled Lapwing, Black Winged Stilt, Pier Avocet, and Common Sandpiper have a strong preference for saline and alkaline lakes that attracts them to SSL. Importantly, some species feeding upon invertebrates little Grebe, Graylag Goose, Bar-Headed Goose, Common Teal, Northern Shoveler, Great Stone Plover, White-Tailed Lapwing, Black-Tailed Godwit, Common Redshank, Curlew Sandpiper, Marsh Sandpiper, Wood Sandpiper, Little Stint, Temmick’s Stint, Ruff, White Wagtail, Grey Wagtail, Pin-tailed Snipe, and Yellow wattled Lapwing found in and around SSL. A species-wise detailed bird census conducted by [12] stated that total 83 waterfowls were recorded. In 1994, 8,500 lesser flamingos were seen on the lake, but no greater flamingoes were found; in 1995, 5,000 lesser flamingoes were recorded but no greater flamingoes were observed, in 2001, 20,000 birds were observed out of which 10,000 were lesser and 5,000 were greater flamingoes.

**Table 1.** Waterbirds counts and comparison.

| S No. | Common and scientific name | 2019 | 2020 |
| --- | --- | --- | --- |
|  | Grebes |  |  |
| 1 | Little Grebe *Tachybaptus ruficollis* | 9 | NF |
|  | HERONS, EGRETS and BITTERNs: |  |  |
| 2 | Black-crowned Night Heron *Nycticorax nycticorax* | 3 | NF |
| 3 | Indian Pond Heron *Ardeola grayii* | NF | 1 |
| 4 | Cattle Egret *Bubulcus ibis* | NF | 5 |
|  | Flamingos: |  |  |
| 5 | Greater Flamingo *Phoeniconaias* | 331 | 12,046 |
| 6 | Lesser Flamingo | 128 | 24,413 |
|  | GEESE and DUCKS: |  |  |
| 7 | Greylag Goose *Anser anser* | 6 | NF |
| 8 | Barheaded Goose *A. indicus* | 18 | NF |
| 9 | Common Pochard *Aythya ferina* | NF | 3 |
| 10 | Gadwall *A. strepera* | 10 | NF |
| 11 | Common Teal *A. crecca* | 9 | NF |
| 12 | Northern Shoveler *A. clypeata* | 359 | 5,293 |
|  | GULLS, TERNS and SKIMMERS: |  |  |
| 13 | Brown-headed Gull *L. brunnicephalus* | NF | 1 |
|  | Plovers: |  |  |
| 14 | Great Stone Plover *Esacus recurvirostris* | 1 | NF |
| 15 | Little-ringed plover *C. dubius* | 4 | 25 |
| 16 | Pacific Golden Plover | NF | 1 |
| 17 | Kentish Plover *C. alexandrinus* | 4 | 47 |
| 18 | Red-wattled Lapwing *V. indicus* | 12 | 16 |
| 19 | White-tailed Lapwing *V. leucurus* | 1 | 2 |
|  | Stilts, avocets: |  |  |
| 20 | Black-winged Stilt *Himantopus himantopus* | 16 | 112 |
| 21 | Pied Avocet *Recurvirostra avosetta* | 34 | 422 |
|  | Snipes, curlews, sandpipers, shanks, godwits, stints: |  |  |
| 22 | Black-tailed Godwit *Limosa limosa* | 2 | 2 |
| 23 | Eurasian Curlew *N. arquata* | NF | 1 |
| 24 | Common Redshank *T. totanus* | 3 | 25 |
| 25 | Common Greenshank *T. nebularia* | NF | 5 |
| 26 | Curlew Sandpiper *C. ferruginea* | 26 | NF |
| 27 | Marsh Sandpiper *T. stagnatilis* | 1 | 2 |
| 28 | Green Sandpiper *T. ochropus* | NF | 2 |
| 29 | Wood Sandpiper *T. glareola* | 5 | 7 |
| 30 | Common Sandpiper *Actitis hypoleucos* | 2 | 15 |
| 31 | Little Stint *C. minuta* | 8 | 110 |
| 32 | Temminck's Stint *C. Temminckii* | 17 | 9 |
| 33 | Ruff *Philomachus pugnax* | 140 | 441 |
|  | Kingfishers: |  |  |
| 34 | White-breasted Kingfisher *H. smyrnens* | NF | 3 |
|  | EAGLES, OSPREY, HARRIERS, FALCONS, KITES: |  |  |
| 35 | Western Marsh-Harrier *Circus aeruginosus* | NF | 1 |
|  | WAGTAILS, PIPIT: |  |  |
| 36 | White Wagtail *Motacilla alba* | 2 | 3 |
| 37 | White-browed Wagtail *M. maderaspatensis* | NF | 1 |
| 38 | Grey Wagtail *M. cinerea* | 1 |  |
|  | Additional species |  |  |
| 39 | Pacific Golden Plover | NF | 1 |
| 40 | Raptor | NF | 3 |
| 41 | Crested Lark | NF | 5 |
| 42 | Greater Coual | NF | 1 |
| 43 | Pintailed Snipe | 5 | NF |
| 44 | Yellow lapwing | 1 | NF |
| 45 | Undefined | NF | 422 |
|  | Total count | 1124 | 43,445 |
|  | Total species no. | 29 | 32 |

## Results

### Past 56 years

Visual interpretation of CORONA revealed four geomorphic units (Aravalli hills, rivers, saline soil, and lake). Two major rivers were identified due to their shape, Mendha in north and Rupnagar in south with their rivulets. Bright tone and smooth textured landform is saline soil, which has reduced (Fig **Error! Reference source not found.**5). Importantly the image of 1972 was classified at 80.95% accuracy with 0.73 Kappa coefficient, 1981 at 82.50 % with 0.76, 1992 at 87.50 % with 0.82, 2009 at 85.71 % with 0.80 and 2019 at 87.50 % at 0.82. This accuracy is reflected in analysis of dynamic matrix.

The study shows degraded trend of LULC for 47 years between 1972-2019 (Table 2, Fig 6). In 1972, the waterbody was 159.6 km^2^ (30%), saltpan was 38.3 km^2^ (7.4%), salt crust was 0 km^2^ (0%), vegetation was 17.9 km^2^ (3.4%), Aravalli was3.5 km^2^ (0.7%), saline soil was 64.3 km^2^ (12.4%) and barren land was 236.0 km^2^ (45.4%). In 1981, waterbody was 98.7 km^2^ (19%), saltpan was 36.1 km^2^ (6.9km2), salt crust was 34.4 km^2^ (6.6%), vegetation as 87.6 km^2^ (16.9%), Aravalli was 3.3 km^2^ (0.6%), saline soil was 49.1 km^2^ (9.4%), barren land was 209.6 km^2^ (40.3%) and settlement was 1.1 km^2^ (0.2%). In 1992, waterbody was 106.7 km^2^ (20.5%), saltpan was 42.8 km^2^ (8.2%), salt crust was 34.7 km^2^ (6.7%), vegetation was 5.3 km^2^ (1.0%), Aravalli was 3.3 km^2^ (0.6%), saline soil was 90.7 km^2^ (17.5%), barren land was 235.3 km^2^ (45.2%) and settlement was 1.1 km^2^ (0.2%). In 2009, waterbody was 31.5 km^2^ (6.1%), saltpan was 64.1 km^2^ (12.3%), salt crust was 0.0 km^2^ (0%), vegetation was 84.1 km^2^ (16.2%), Aravalli was 3.2 km^2^ (0.6%), saline soil was 118.3 km^2^ (27.7%), barren land was 217.3 km^2^ (41.8%) and settlement was 1.4 km^2^ (0.3%). In 2019, waterbody was 17.4 km^2^ (3.4%), saltpan was 72.9 km^2^ (14.0%), salt crust was 15.4 km^2^ (3.0), vegetation was 34.1 km^2^ (6.6%), Aravalli was 3.2 km^2^ (0.6%), saline soil was 112.6 km^2^ (21.7%), barren land was 257.8 km^2^ (49.6%) and settlement was 6.5 km^2^ (1.3%). Overall, the change from 1972 to 2019 has been summarized, as waterbody decreased from 30.7 to 3.4%. Salt crust increased from 0 to 3%. Vegetation increased from 3.4 to 6.6%. Aravalli decreased from 0.7 to 0.6%. Saline soil increased from 12.4 to 21.7%. Barren land increased from 45.4 to 49.6%. Saltpan increased from 7.4 to 14%. Settlement increased from 0.1 to 1.3%.

**Table 2.**
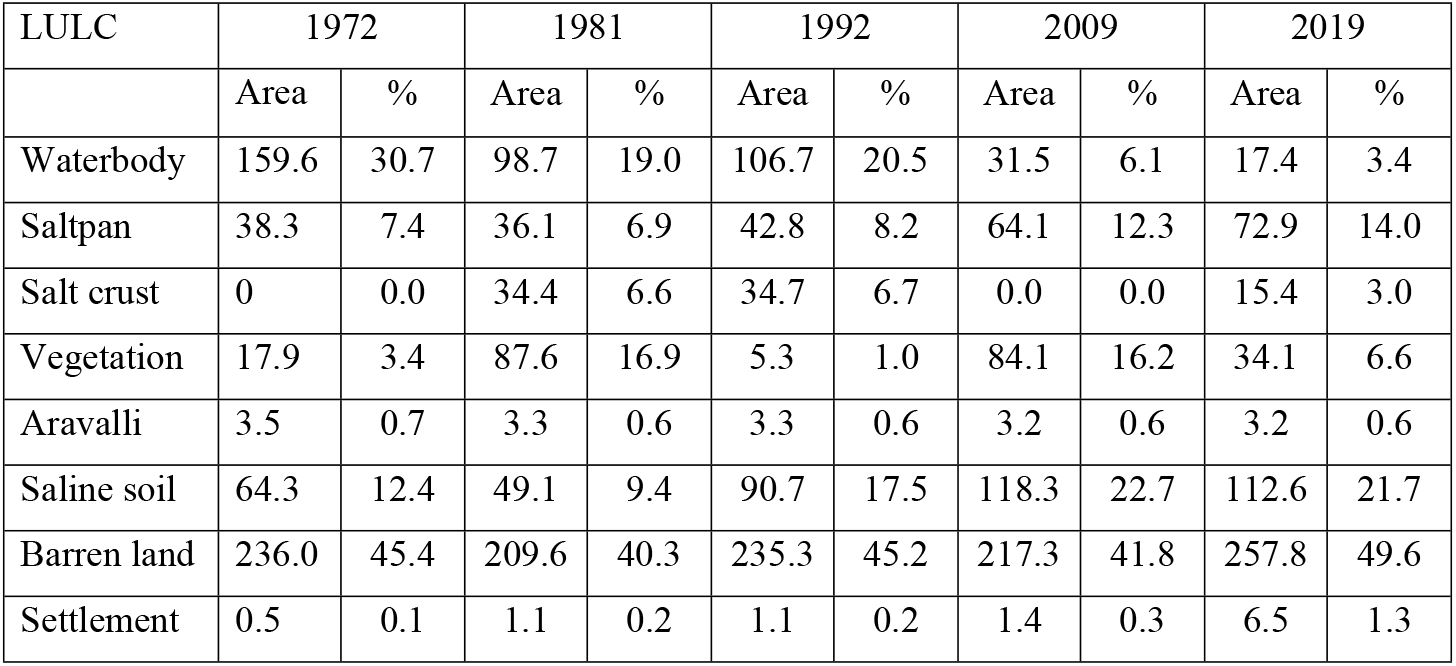
LULC change area from 1972-2019 (area in km^2^).

**Fig 6.**
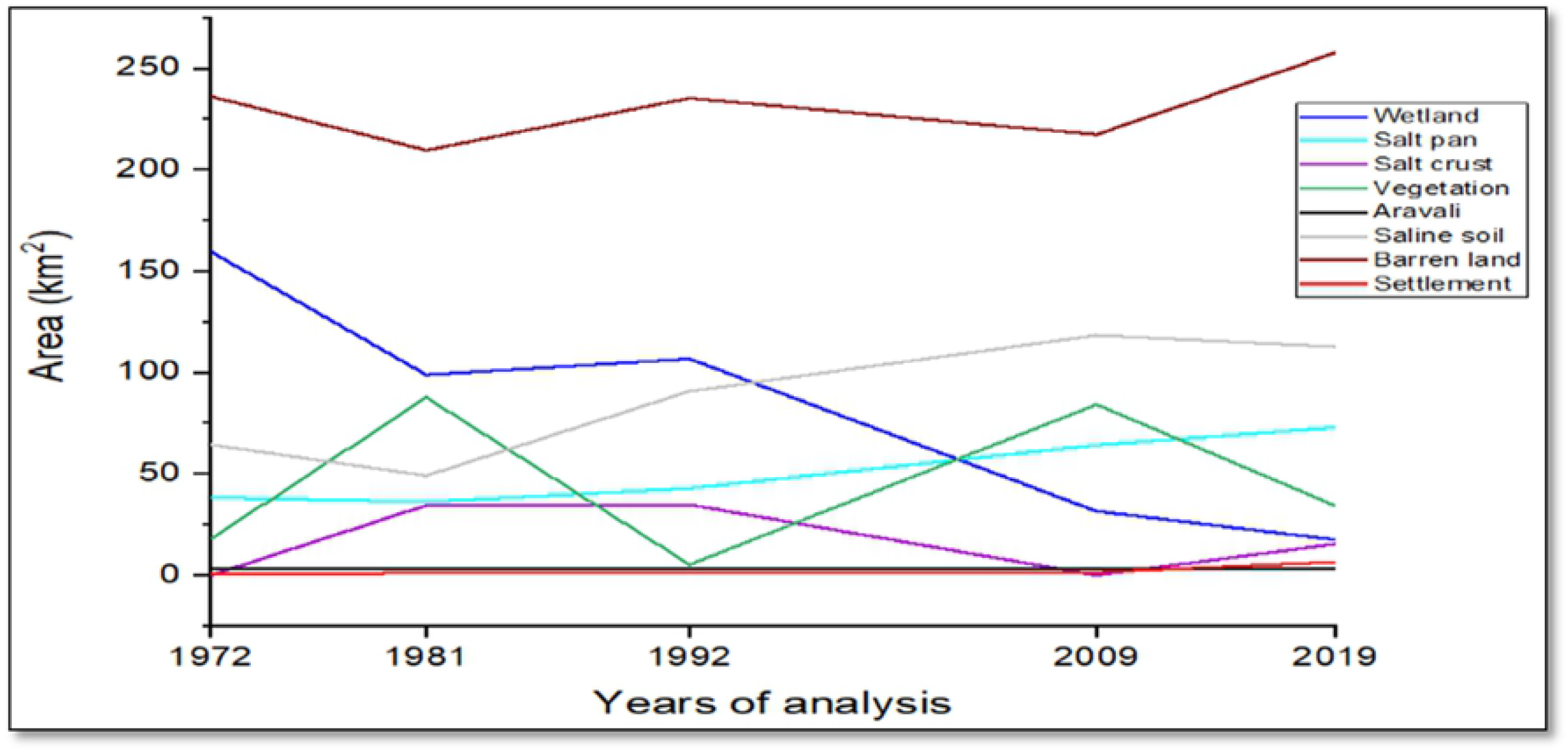
Trend of LULC change.

LULC change rate (Table 3) is represented as K (%). It shows the waterbody degrading at the rate of −4.23%, 0.73%, −4.14%, and −4.47% since 1972-2019. In fact, in the first decade, K of vegetation was 43.38% and settlement was 12.66%. Saltpan decreased by 0.63%, Aravalli by 0.56%, saline soil by 2.62%, and barren land by 1.24%. Furthermore, from 1981 to 1992, only vegetation changed negatively; rest of the classes like increased wetland by 0.73%, saltpan by 1.68%, salt crust by 0.08%, Aravalli by 0.06%, saline soil by 7.71%, barren land by 1.11% and settlement by 0.43%. From 1992 to 2009, wetland decreased by −4.14% followed by salt crust by 5.88%, Aravalli by 0.11%, and barren land by 0.45% whereas vegetation by 0.20%, and saline soil by 1.78% positive K. From 2009 to 2019, wetland, vegetation, Aravalli, salt crust, and saline soil showed negative K by 4.47%, 5.95%, 0.11%, 0.00%, and 0.48% respectively and saltpan, barren land, and settlement showed positive K of 1.36%, 1.86%, and 37.98% respectively. Settlement has high K value in this decade.

**Table 3.** LULC dynamic degree K (percentage).

| LULC classes | 1972-81 | 1981-92 | 1992-09 | 2009-19 |
| --- | --- | --- | --- | --- |
| waterbody | -4.23% | 0.73% | -4.14% | -4.47% |
| Saltpan | -0.63% | 1.68% | 2.93% | 1.36% |
| Salt crust | 0.00% | 0.08% | -5.88% | 0.00% |
| Vegetation | 43.38% | -8.54% | 88.20% | -5.95% |
| Aravalli | -0.56% | 0.06% | -0.11% | -0.11% |
| Saline soil | -2.62% | 7.71% | 1.78% | -0.48% |
| Barren land | -1.24% | 1.11% | -0.45% | 1.86% |
| Settlement | 12.66% | 0.43% | 1.25% | 37.98% |

LULC transition matrix (Table 4) states conversion from wetland (75 km^2^) to saline soil, second largest from barren land to (22.5 km^2^) to vegetation. 0.96 km^2^ of Aravalli to barren land, saline soil (21.67 km^2^) to barren land. 13.87 km^2^ and 12.11 km^2^ of salt crust is to saline soil and barren land respectively.

**Table 4.** LULC transition matrix from 1981-2019.

| 1981 | 2019 |  |  |  |  |  |  |  |  |
| --- | --- | --- | --- | --- | --- | --- | --- | --- | --- |
|  | Aravalli | Barren land | Saline soil | Salt crust | Saltpan | Settlement | Vegetation | Water body | Grand Total |
| Aravalli | 2.12 | 0.96 | 0.00 | 0.00 | 0.01 | 0.01 | 0.11 | 0.01 | 3.23 |
| Barren land | 1.02 | 155.46 | 3.54 | 1.04 | 21.26 | 3.25 | 22.57 | 0.55 | 208.69 |
| Saline soil | 0.00 | 21.67 | 17.86 | 2.96 | 4.47 | 0.04 | 1.97 | 0.10 | 49.08 |
| Salt crust | 0.00 | 12.11 | 13.87 | 4.33 | 2.99 | 0.10 | 0.95 | 0.08 | 34.43 |
| Saltpan | 0.00 | 0.89 | 0.32 | 1.92 | 29.21 | 0.33 | 0.96 | 2.41 | 36.04 |
| Settlement | 0.08 | 66.07 | 1.22 | 0.26 | 11.63 | 2.78 | 7.35 | 0.40 | 89.79 |
| Wetland | 0.00 | 0.70 | 75.84 | 4.86 | 3.30 | 0.02 | 0.16 | 13.87 | 98.74 |
| Grand Total | 3.22 | 257.85 | 112.65 | 15.38 | 72.86 | 6.52 | 34.09 | 17.43 | 520.01 |

### Current status of SSL

#### Soil and water quality parameters

Soil parameters like pH, Electrical Conductivity (EC), salinity, chloride, carbonate, and Total Organic Carbon (TOC) were analyzed and mapped (Fig 7). Linear correlation was calculated. Highest positive correlation was observed between salinity and EC (r = 0.99), indicating salts as the major factors for conductivity. Anions like chloride, carbonates, and bicarbonates, chloride have very high EC (r = 0.99), which infers that chlorides are major anions responsible for salinity. TOC is slightly positively related (r =0.5) to other parameters. Water parameters like pH, EC, TDS, salinity, chloride, carbonate, and hardness were calculated and mapped (Fig 8). Linear correlation was calculated. Highest positive correlation was observed between salinity and EC (r=0.93) and the same positive relationship between salinity and TDS. As per the analysis, major salinity has been contributed by chloride ions; it also shows positive correlation between EC and TDS. Other than chloride, carbonate has also correlation with TDS (r=0.5).

**Fig 7.**
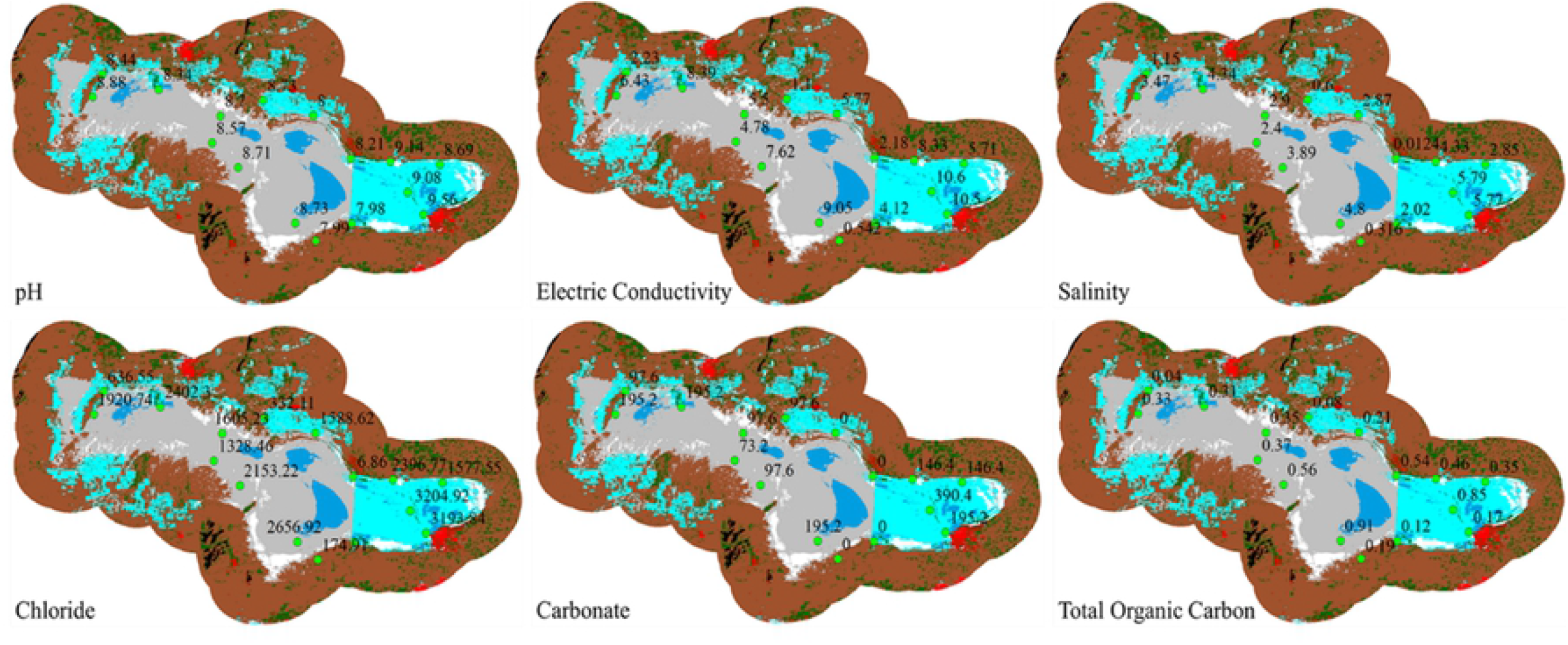
Mapping soil quality parameters.

**Fig 8.**
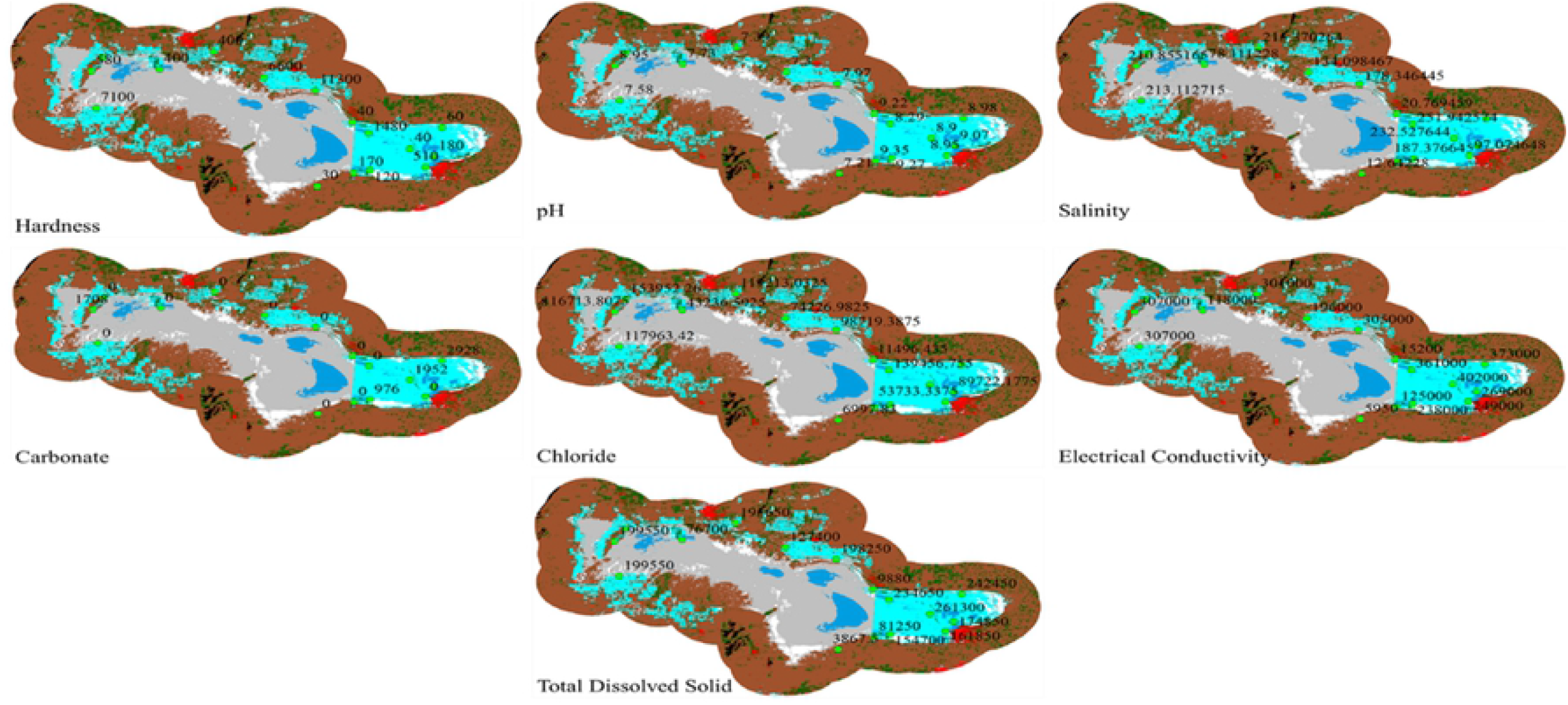
Mapping water quality parameters.

### Future LULC for next 40 years

Future prediction was conducted. Predicted maps of 2029, 20239, 2049, and 2059 were obtained (Fig 9). Waterbody has been interchangeably used with wetland class name. There will be decrease in wetland area and increases of saltpans towards north. Conversion of saline soil into barren land in central part is observed. Towards south, saltpans will have no noticeable change until 2039, however, will increase in 2049 and 2059 maps. Area statistics graphs for predicted maps were derived through modelling (Fig 9a-j). (a) shows percentage wise gain and loss; (b) shows net change in percentage and (c to j) shows class-wise contributions to net changes.

**Fig 9.**
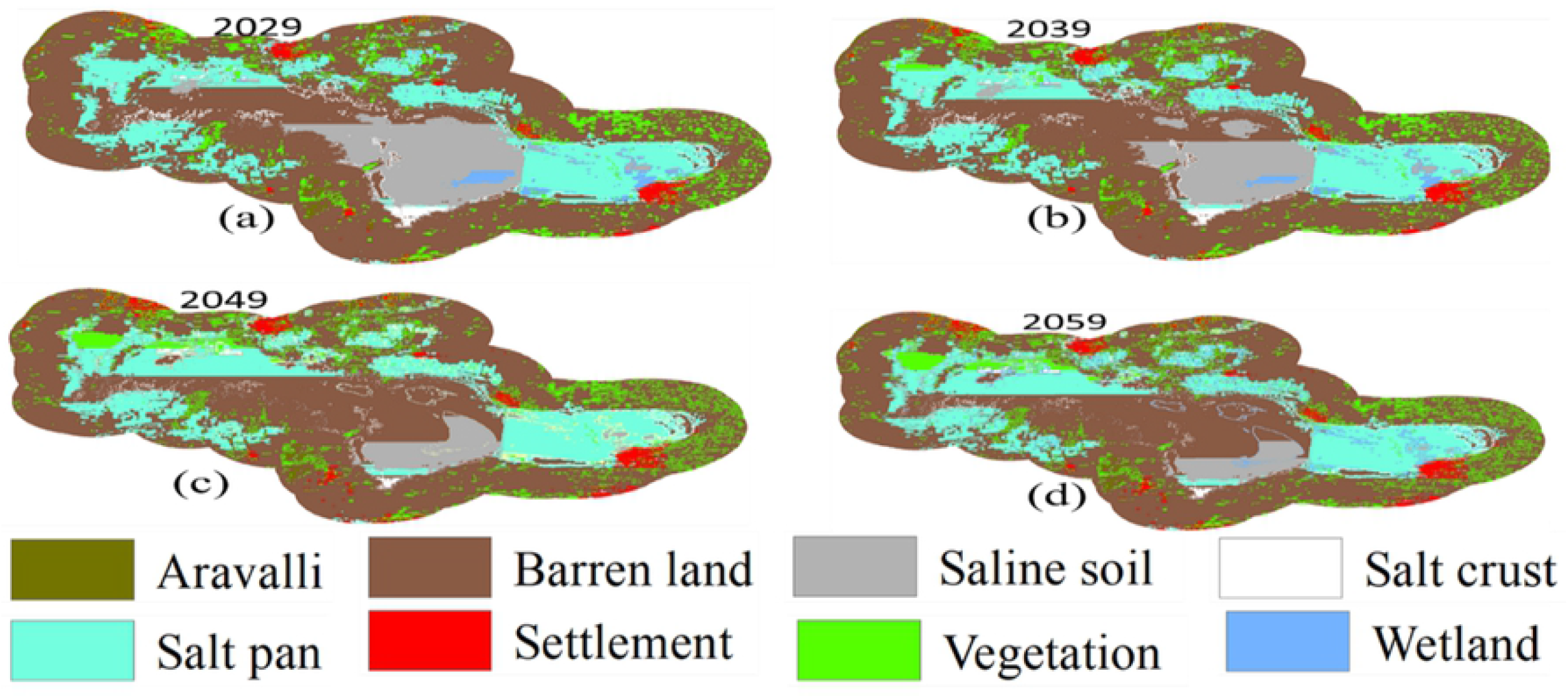
Maps of future prediction.

Fig 10 (a) shows highest gain in wetland by 60% and highest loss of saline soil by 70%. It shows wetland loss by 40%, vegetation gained 40% and loss 30%, settlement gained by 50% and loss by 20%, saltpan gain by 30% and loss by 20%, salt crust gain by 40%, and loss by 50%, saline soil gain by 40% and loss by 70%, barren land gain by 20% and loss by 10 % and Aravalli gain by 40% and loss by 20%. Fig 10 (b) shows net increase by 20% in wetland, 30% in vegetation, 40% in settlement, 10% in saltpan, 5% in barren land, and decrease by 20% in salt crust, saline soil by 120%, and Aravalli by 20%. Fig 10 (c to j) shows net change in each LULC. Aravalli will positively impact by 0.01% and negatively by 0.02% and 0.04% by settlement and barren land. Positive contributions to net change of barren land by salt crust and saline soil is by 0.01% and 5% respectively, negatively in wetland by 0.01%, vegetation by 0.5%, settlement by 0.05%, and saltpan by 3%. Positive contributions to saline soil are by wetland and saltpan by 0.1% and 0.5% and negatively by 5% by barren land. Positive contribution in salt crust by 0.1% and negatively by vegetation and barren land by 0.2% and 0.24% respectively. Positive contributions to saltpan by barren land is by 2.79% and negatively by wetland, vegetation, and saline soil. Settlement would experience positively by vegetation, saltpan, barren land, and Aravalli by 0.18, 0.02, 0.34, and 0.1 % respectively and negatively by saline soil by 0.01 %. Vegetation will experience positively by saltpan by 0.8 % and barren land by 0.7 % and negatively by settlement by 0.21 % and lastly wetland will experience positively by saltpan by 0.30 %, barren land by 0.10 % and negatively by saline soil by 0.25 %. Spatial cubic trend changes of SSL are given in (Fig 11).

**Fig 10.**
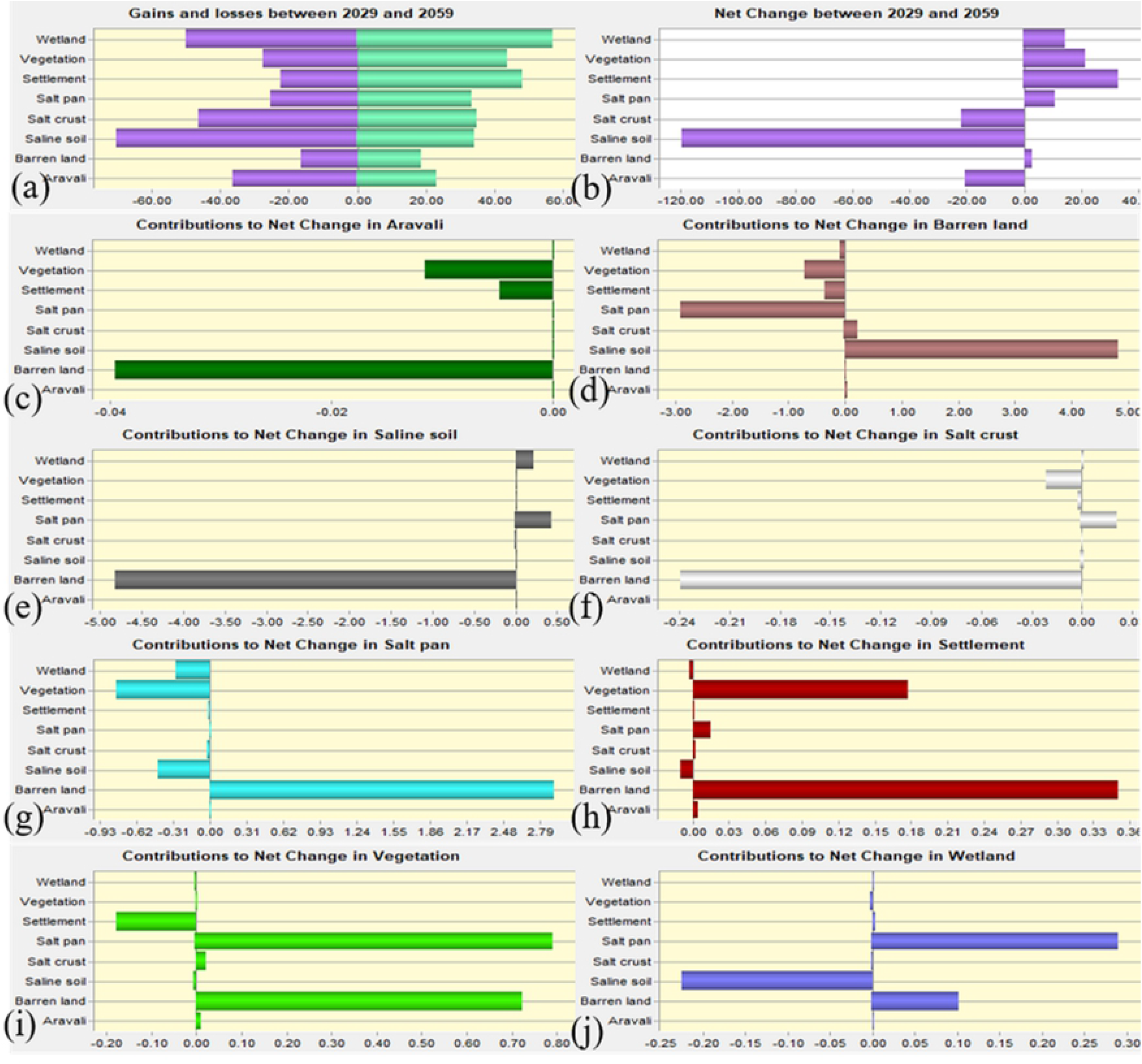
LULC change percentages: (a) gain and loss of LULC (b) net change between 2029 and 2059 and (c-j) contributions of each class to net changes.

**Fig 11.**
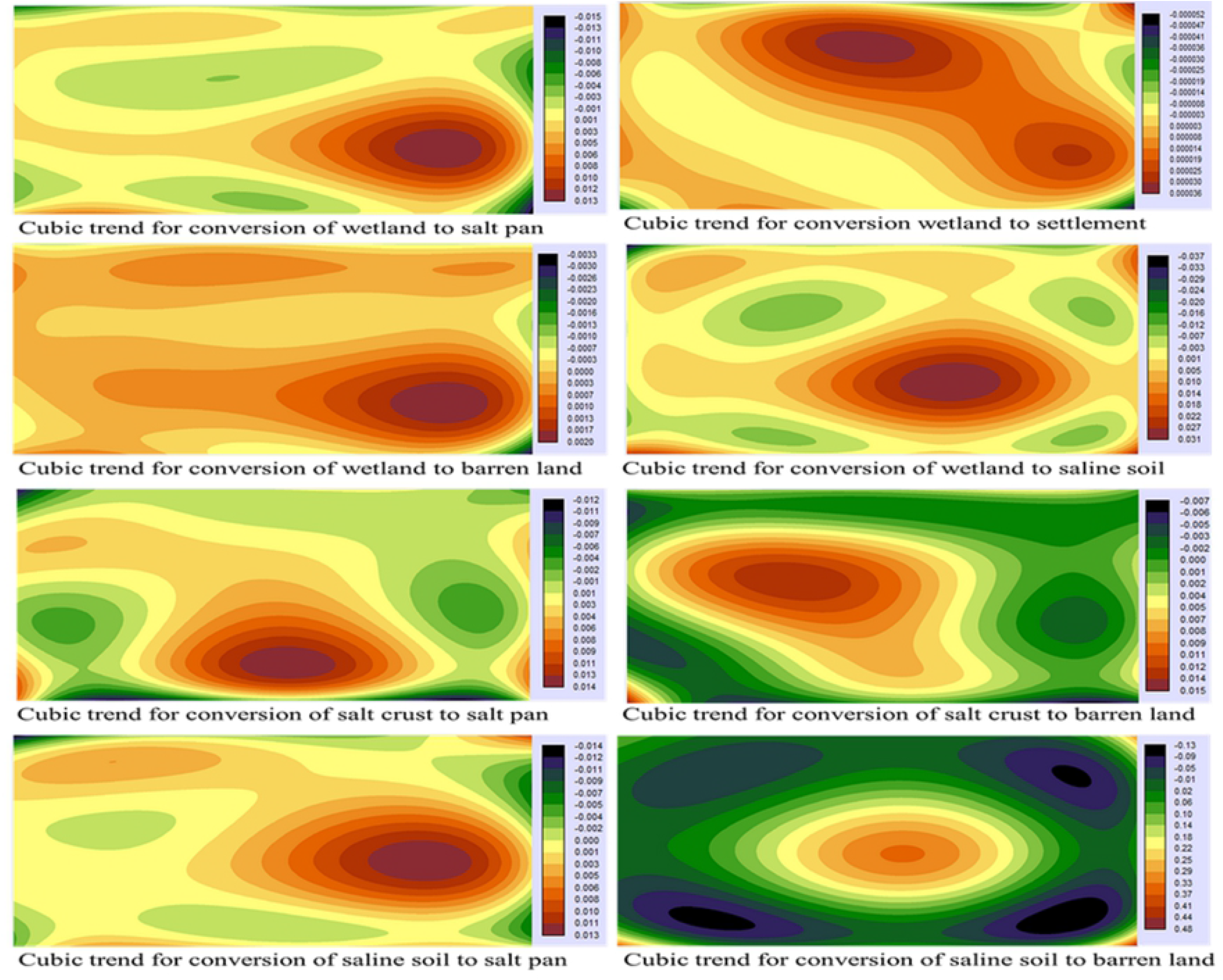
Spatial trend representation.

## Discussion

### Loss of saline character

The topmost layer of SSL (0-115 cm) appears greyish brown to dark grey from top to bottom due to organic residues with minerals of Kieserite providing brine rich in carbonate, sulfate, calcium, magnesium, 115-360 cm appear dark grey with bloedite and brine rich in sodium, 360-600 cm is rich in calcite, polyhalite, gypsum, dolomite and brine rich in same constituents besides potassium, 600-1900 cm has weathered gypsum, calcite, dolomite, polyhalite, thenardite with no brine and below 1900 cm has pre-Cambrian rock basement consisting of schists, phyllites and quartzite [10]. However, the decadal analysis states that the six vertical soil gradients are at stake. The only method approved by GoI is the collection of brine through pans and kyars [41]. 18,65,000 tons/year of salt were sustainably extracted in which lake water provided 18*105 tons/ year, rainwater 5*103 tons/year, and river water 6*104 tons/ year [11]. 2 000 illegal tube wells and 240 bore wells have been built over the last two decades. [42] stealing brine worth 300 million USD [41] from both surface and sub-surface [10]. Results from the imagery of 1963 do not suggest that there were unauthorized pans. It occupied 7.4 % by authorized Sambhar Salt Ltd. in 1972. Gradually in 1992, 2009 and 2019 encroachment increased to 8.2, 12.3 to 14, 10 % respectively. Major encroachment appeared in Nagaur due to the construction of hydrological structures [43]. Other threats are livestock ranching, poaching, sewage discharge, trails [36], vehicle testing [42].

### Loss of wetland connectivity and trophic structure

Within the 3 km SSL buffer zone, Naliasar, Devyani sarovar, and Ratan talab are linked by birds for breeding, feeding, and roosting. Their connectivity depends on water budget, hydrophytes, hydric soil, predator status, food availability, hydro-period, wetland complexes, topography, geography, and weather [44]. However, the results of satellites show a steady decline at 4 % from 30.4 to 3.4 %. This has forced the bird to move elsewhere. Due to the shrinkage, its complex trophic structure with 39 aquatic and 80 terrestrial producers, 133 primary and secondary consumers [25], and 279 birds as tertiary consumers [12] are at stake. Depending on water availability, level, depth, and microbiota, wetland connectivity is divided into three types; include bottom-dwelling, surface, and shore animals for SSL [23]. Bottom and mud dwelling animals like *Polypodium sp.* and *Chironomus sp.* survive in favorable seasons from July to December, when salinity is 9.6 to 72.6 %, carbon dioxide is in between 48 to 56.2 mg/l, and oxygen is between 42 to 27.8 mg/l [23]. Surface animals consist of both plankton and nekton. Phytoplanktons *(Dunaliella saline, Aphanotheca halophytica, Spirulina sp)* and zooplanktons protozoans, nauplii of crustaceans [23], and nektons are stenohaline that survive during the favorable condition, and replaced by euryhaline animals *(Artemia salina, Ephydra macellaria,* and *Eristalis sp.)* during adverse conditions, tolerating up to 164 % salinity, and disappear in May to June, when lake is naturally dry. Shore animals represented by *Labidura riparia, Coniocleonus sp.* and others survive during favorable periods; however, they travel to the core during adverse conditions. However, these species might not be available as the lake is shrinking.

### Management and Restoration potentials

When saline lakes are relentlessly desiccated, they might become dust bowl harmful for both man and environment as in case of Owens Lake in California or collapse billion-dollar global market of brine shrimp as in case of Lake Urmia or loss of 40,000 metric tons of fishery and 60,000 jobs in Aral Sea [45]. Shriveled saline lakes create ecological disconnectivity, neither support unique halophytes and halophytes nor attract flamingoes or other birds [46]. So, this is the high time when Sambhar lake requires urgent attention, least it might also require more capital for restoration than it generated revenue as in the case of Owen’s lake when US$ 3.6 billion was spent for its dust mitigation [47]. At current stage, it is possible through the reconstruction of its physicochemical adjustments, and reintroduction of native flora and fauna. Emphasis on health protection, incentives, and rewards be given to salt workers so that more people participate in wise use of this lake. Demolishing check dams and anicuts, ban of sub-surface brine collection, using electrical pumps, illegal salt pan encroachment in and around lake be declared as a punishable act, demolish construction up to 3km buffer zone declaring it ‘no construction zone’, controlled sewage disposal, increasing water residence period. Increasing aquatic biodiversity, hydrodynamics, nutrient cycling, vegetative and non-vegetative productivity, cascading trophic levels be focused. These steps will not only help SSL to its pristine conditions but also generate revenue for a longer period, provide jobs to more people and attract more migratory birds.

### Conclusion

This work emphasizes the understanding of spatio-temporal variation of the largest inland saline Ramsar site in semi-arid region of India over 9 decades. Past LULC trends indicated its continuous shrinkage. This study further confirmed the current situation through field visits, soil-water parameters analysis, bird census information. We modelled its future predictions using the CA-Markov model using geospatial platform until 2059. It indicated complete desiccation of the wetland to a wasteland. This lake will neither be able to generate billion dollars revenue, nor attract lakhs of migratory birds or provide jobs to thousands of salt workers. The major influencing factors are illegal saltpan-encroachment, excess groundwater extraction, increasing settlement area, and water diversion. Our study suggests that its restoration is very much essential to revive its hydrological configurations. Further, we have also suggested some restoration strategies that could be practically implemented.

## Author contributions

Both the authors contributed to the study conception, design, and data collection. Methodology was conceptualized by Sharma.L and was performed by Naik.R. The first draft of the manuscript was written by Naik.R. Sharma.L provided editorial comments. Both the authors read and approved the final manuscript.

